# Prime-boost protein subunit vaccines against SARS-CoV-2 are highly immunogenic in mice and macaques

**DOI:** 10.1101/2020.09.01.278630

**Authors:** Hyon-Xhi Tan, Jennifer A Juno, Wen Shi Lee, Isaac Barber-Axthelm, Hannah G Kelly, Kathleen M Wragg, Robyn Esterbauer, Thakshila Amarasena, Francesca L Mordant, Kanta Subbarao, Stephen J Kent, Adam K Wheatley

## Abstract

SARS-CoV-2 vaccines are advancing into human clinical trials, with emphasis on eliciting high titres of neutralising antibodies against the viral spike (S). However, the merits of broadly targeting S versus focusing antibody onto the smaller receptor binding domain (RBD) are unclear. Here we assessed prototypic S and RBD subunit vaccines in homologous or heterologous prime-boost regimens in mice and non-human primates. We find S is highly immunogenic in mice, while the comparatively poor immunogenicity of RBD was associated with limiting germinal centre and T follicular helper cell activity. Boosting S-primed mice with either S or RBD significantly augmented neutralising titres, with RBD-focussing driving moderate improvement in serum neutralisation. In contrast, both S and RBD vaccines were comparably immunogenic in macaques, eliciting serological neutralising activity that generally exceed levels in convalescent humans. These studies confirm recombinant S proteins as promising vaccine candidates and highlight multiple pathways to achieving potent serological neutralisation.

## Introduction

The rapid onset and global spread of the SARS-CoV-2 pandemic has spurred unprecedented global scientific efforts to develop, test and manufacture novel protective vaccines. The spike (S) glycoprotein of SARS-CoV-2 is a clear target for vaccines designed to elicit neutralising antibodies to prevent infection. Recent studies suggest neutralising antibodies can protect macaques against SARS-CoV-2 (Deng et al., 2020; Guebre-Xabier et al., 2020; Zost et al., 2020) and observational human studies also suggest neutralising antibody responses are protective against re-infection (Addetia et al., 2020). S is a type 1 viral fusion protein, expressed as a single polypeptide and cleaved into S1 and S2 subunits, with a hetero-trimeric quaternary structure common to many respiratory viruses (reviewed in Li, 2016). Cell entry is mediated by engagement of the enzyme ACE2 on the target cell surface by the viral receptor binding domain (RBD), localised within the C-terminal domain of S1 (Lan et al., 2020; Walls et al., 2020). Antibodies capable of preventing RBD binding to ACE2 can therefore prevent infection and constitute an efficient pathway to neutralisation.

Viruses employ numerous strategies to avoid immune recognition of viral entry proteins, including heavy decoration with N-linked glycans (Watanabe et al., 2020), employing immune distraction or escape by focussing host immunity onto highly mutable regions (Li et al., 2020). A shared challenge for vaccine development against viral glycoproteins is therefore ensuring maximum B cell recognition of neutralising epitopes critical for viral replication (on-target), while minimising responses to poorly conserved epitopes or those with no antiviral capacity (off-target).

Currently, several S-based vaccines are entering early stage clinical trials, including recombinant trimeric S proteins, monomeric or trimeric RBD domains, and analogues delivered by viral vectors or mRNA (Folegatti et al., 2020; Jackson et al., 2020; Keech et al., 2020; Mulligan et al., 2020; Walsh et al., 2020; Zhu et al., 2020). However, the relative merits of these immunogens are currently unclear. In particular, does the use of a smaller vaccine target, such as the RBD, drive a more focussed neutralising antibody response? Or alternatively, do additional epitopes across the larger S protein make additive contributions to immunogenicity or neutralisation that counteract any off-target immune distraction? Here we directly compare the immunogenic profile of SARS-CoV-2 S and RBD immunogens in mice and non-human primates using various prime-boost approaches. We find in mice that RBD is relatively poorly immunogenic compared to S, with primary immunisation compromised by a reduced capacity to efficiently induce germinal centre B cells and recruit effective T follicular helper cells. In contrast, immunisation with S alone, or boosting S-primed animals with S or RBD, is potently immunogenic, reliably eliciting strong binding and neutralising titres in immunised mice. In more genetically diverse non-human primates, two immunisations with either S or RBD immunogens were comparably immunogenic and produced strong serological neutralising responses. Overall, we find that immunisation with recombinant S immunogens reliably elicits potentially protective humoral immunity at levels in excess of those observed in convalescent humans.

## Results

### SARS-CoV-2 spike but not RBD is potently immunogenic in mice

The primary immunogenicity of SARS-CoV-2 S and RBD was assessed in groups of C57BL/6 mice vaccinated with S, RBD or control ovalbumin (OVA) proteins. A single immunisation of S formulated with the adjuvant Addavax was highly immunogenic, eliciting high reciprocal serum endpoint titres of S-specific antibody (Figure 1A; median 1.85 x 10^5^; IQR 1.34-4.61), but without inducing significant neutralisation activity (Figure 1B). In contrast, a single immunisation of RBD elicited minimal serum antibody in line with other reports of sub-optimal immunogenicity of RBD in mice (Mandolesi et al., 2020) and recent Phase I trials comparing RBD or S encoded by RNA-based vaccines (Walsh et al., 2020). Although S is extensively glycosylated (Watanabe et al., 2020), limiting complex glycan deposition by expression in HEK293 cell lines lacking N-acetylglucosaminyltransferase I (S gnt-) did not negatively impact the potent immunogenicity of S (Figure 1A).

**Figure 1.**
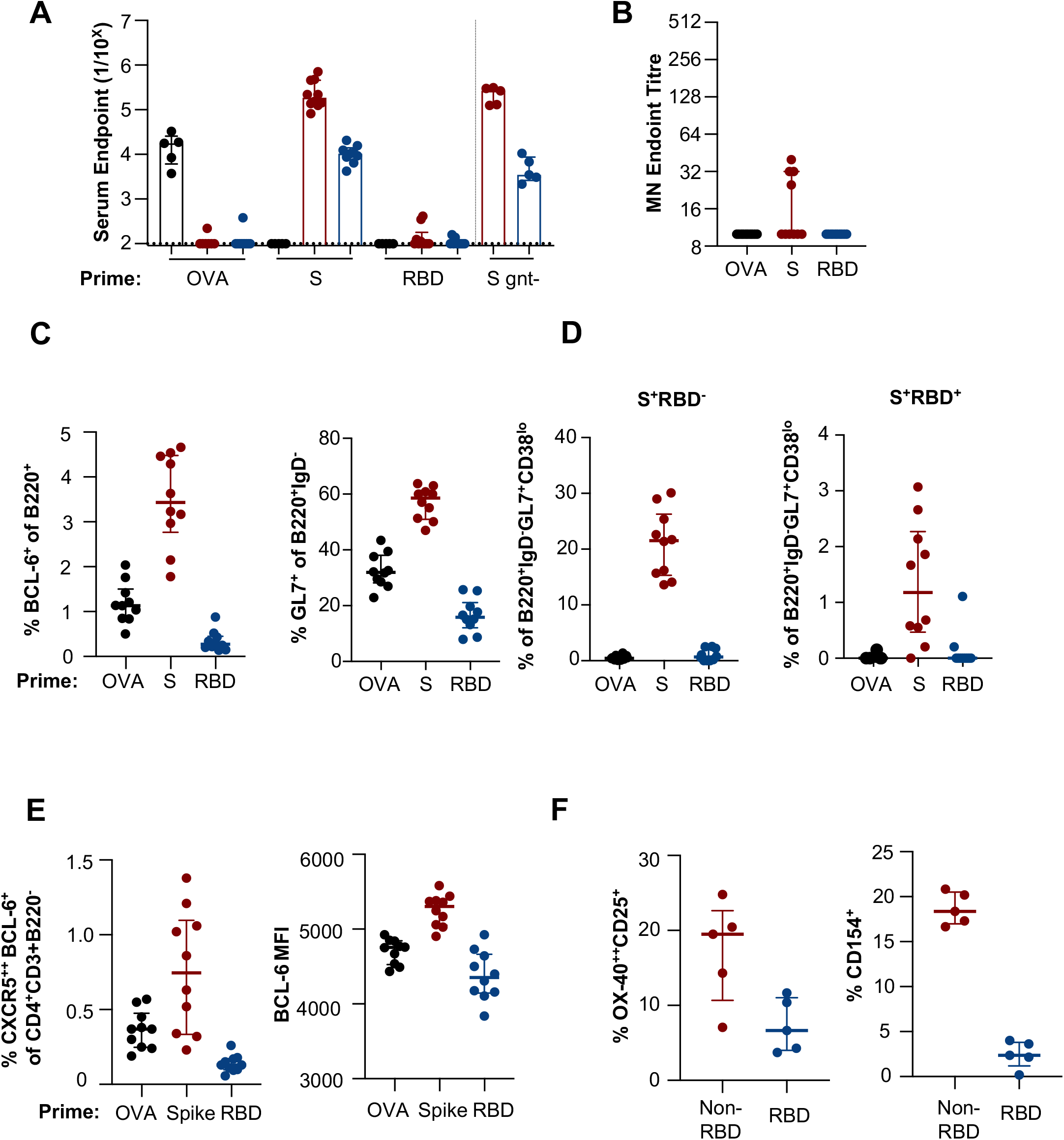
Primary immunogenicity of SARS-CoV-2 subunit proteins in C57BL/6 mice. Mice were immunised intramuscularly with S, RBD or OVA proteins and immune responses assessed 14 days post-immunisation (n = 10, S gnt-n = 5). **(A)** Reciprocal serum endpoint dilutions of S-(red), RBD-(blue) or OVA-specific IgG (black) were measured by ELISA. Dotted lines denote the detection cut off (1:100 dilution). **(B)** Serum neutralisation activity was assessed using a microneutralisation assay. **(C)** Draining lymph node germinal centre activity assessed by BCL-6 expression in B220^+^ B cells or GL7 expression in B220^+^IgD B cells. **(D)** Frequency of germinal centre B cells (B220^+^IgD GL7^+^CD38^lo^) specific for Spike (S^+^RBD) or RBD (S^+^RBD^+^) probes. **(E)** Frequency of TFH cells (CXCR5 BCL-6^+^CD4^+^CD3^+^B220) and corresponding median fluorescent intensity of BCL-6. **(F)** Antigen-specific TFH cells were identified as either OX-40^++^CD25^+^ or CD154^+^ following 18 hours of stimulation with S (minus RBD) or RBD peptide pools. Antigen-specific responses are presented after background subtraction using a DMSO control. Data is presented as median ± IQR.

The elicitation of SARS-CoV-2 specific B and T cell responses in the draining iliac and inguinal lymph nodes (LN) was assessed by flow cytometry. S-immunised animals displayed robust induction of germinal centre (GC) B cells measured either by intracellular BCL-6^+^ (median 3.4%; IQR 2.8-4.5) or surface GL7^+^ (median 58.5%; IQR 51.0-63.8) expression (Figure 1C; gating in Figure S1). Immunisation with OVA induced intermediate frequencies of BCL-6^+^ (median 1.1%; IQR 0.8-1.5) and GL7^+^ (median 32.0%; IQR 28.2-38.1) B cells, with minimal detection of GC formation in RBD-immunised animals (BCL-6^+^; median 0.3%; IQR 0.2-0.4) (GL7^+^; median 15.9%; IQR 12.1-21.1). The specificity of GC B cells (IgD^-^B220^+^GL7^+^CD38^lo^) was examined using recombinant S and RBD probes, as previously described (Juno et al., 2020). Both S-specific (S^+^RBD^-^) and RBD-specific (S^+^RBD^+^) were reliably detected in S-immunised animals, constituting 21.5% (IQR 15.3-26.3) and 1.2% (IQR 0.47-2.27) of GC B cells respectively (Figure 1D). Mirroring the serum antibody response, few S- or RBD-specific GC B cells were observed in RBD-immunised animals.

Consistent with GC B cell frequencies, immunisation with S induced high frequencies of T follicular helper (TFH) cells (CXCR5^++^BCL-6^+^; median 0.7%; 0.3-1.1) relative to OVA (median 0.4%; 0.2-0.8) or RBD (median 0.13%, 0.1-0.2) (Figure 1E; gating Figure S2). The median fluorescent intensity (MFI) of BCL-6 expression in TFH was notably higher in S-vaccinated animals (median 5303 for S, 4756 for OVA and 4353 for RBD) (Figure 1E). Analysis of TFH specificity using an activation induced marker (AIM) assay (Jiang et al., 2019) and pools of RBD or non-RBD S overlapping peptides indicated that the TFH response is dominated by non-RBD-localised epitopes in S-immunised animals (Figure 1F). Further, RBD peptides elicited weak CD154 responses upon re-stimulation in comparison to non-RBD peptides (Figure 1F). Epitope mapping of peptides spanning the RBD indicated that only 3 RBD-derived peptides were recognised by CD4 T cells in C57BL/6 mice (Figure S3), one of which was substantially more immunogenic than the others (P99, sequence TNVYADSFVIRGDEV). In contrast, a more extensive number (>8) of S epitopes were identified outside the RBD (4 of which are shown in Figure S3). In S-vaccinated animals, the 3 RBD-derived epitopes were subdominant to the most immunogenic non-RBD S epitopes and relatively poor inducers of CD154 expression (Figure S3). These data suggest that the presence of the single relatively immunogenic TFH epitope (P99) within the SARS-CoV-2 RBD is insufficient to drive a robust CD4 T cell response in comparison to the greater number of immunodominant epitopes available in full-length S.

The primary immunogenicity of the RBD was similarly muted in BALB/c mice in comparison to S, with poor induction of RBD-directed antibody, GC B cells and TFH responses (Figure S4). Overall, we find that S is potently immunogenic in both mouse strains, with GC B cell and TFH responses largely targeted at regions outside the RBD. In contrast, the immunogenicity of the RBD is constrained in mice, likely in part through suboptimal recruitment of quality TFH responses.

### Homologous spike and heterologous spike/RBD prime-boost immunisations elicit potent binding and neutralising antibody responses in mice

Despite potentially compromised immunogenicity, the small antigenic target of the RBD remains attractive for focusing immunity upon protective neutralising epitopes. However, the recent identification of N-terminal domain (NTD) (Chi et al., 2020) and other alternative S-localised protective epitopes (Liu et al., 2020; Rogers et al., 2020) highlights additional antibody targets for vaccine protection. To assess the relative merits of RBD-focussed antibody responses versus more holistically targeting the entire S, we primed C57BL/6 mice with S or RBD, and then boosted 21 days later with either homologous or heterologous immunogens. Homologous S prime-boost (S-S) elicited high reciprocal serum endpoint titres of both S- (median 2.77 x 10^6^; IQR 1.71-3.62) and RBD-specific antibodies (2.28 x 10^6^; IQR 1.23-3.17) (Figure 2A). In contrast to a single RBD dose, we found that homologous RBD prime-boost (R-R) immunisation was capable of eliciting modest serum titres of S- and RBD-specific antibodies. RBD prime-S boost (R-S) induced similarly modest titres. However, we find RBD boosting of S-primed animals (S-R) markedly increased RBD-specific serum antibody titres 4.2-fold relative to S-S (p=0.0055). Inhibition of ACE2-RBD interaction by serum antibodies was similarly enhanced in the S-R group compared to the S-S group (Figure 2B, Figure S5). Selective RBD focussing translated into increased serum neutralisation, with S-R eliciting 2.5-fold higher activity (median 361; IQR 226-706) compared to S-S (143; IQR 96-254) (p=0.0055) (Figure 2C). Neither R-R nor R-S immunisation reliably induced significant serum neutralising activity, although they elicited modest levels of antibodies that block ACE2-RBD engagement. The capacity of S-S and S-R immunisation to elicit potent binding and neutralising responses in C57BL/6 mice was mirrored in BALB/c mice (Figure S6).

**Figure 2.**
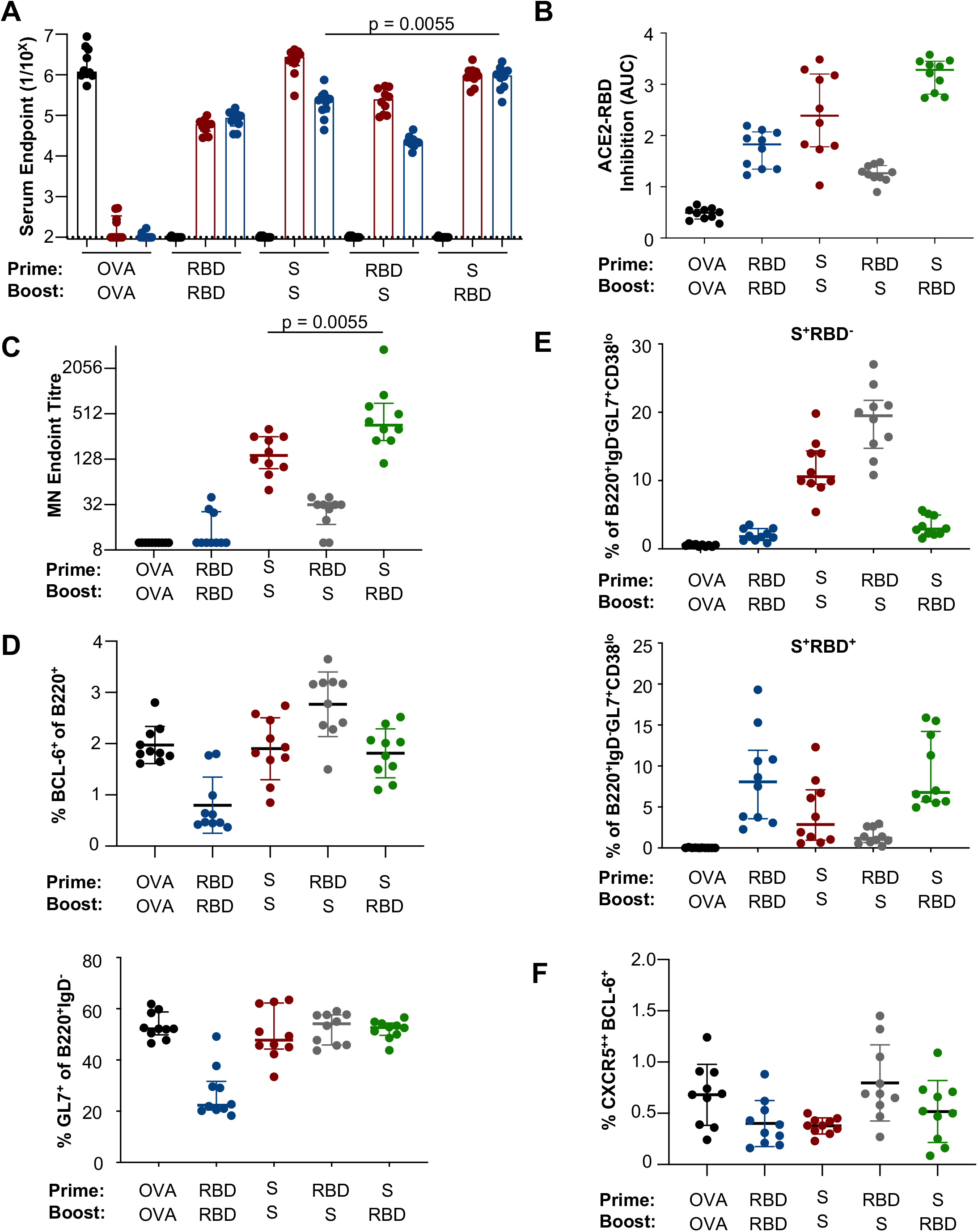
Prime-boost immunisation of SARS-CoV-2 subunit proteins in C57BL/6 mice. Mice were serially immunised intramuscularly at a 21-day interval with S, RBD or OVA proteins and immune responses assessed 14 days post-boost (n = 10). **(A)** Reciprocal serum endpoint dilutions of S-(red), RBD-(blue) or OVA-specific IgG (black) were measured by ELISA. Dotted lines denote the detection cut off (1:100 dilution). **(B)** The capacity of serum antibodies to inhibit the interaction of RBD and human ACE2 was assessed by ELISA. **(C)** Neutralisation activity in the serum was assessed using a microneutralisation assay. **(D)** Draining lymph node germinal centre activity assessed by BCL-6 expression in B220^+^ B cells or GL7 expression in B220^+^IgD B cells. **(E)** Frequency of germinal centre B cells (B220^+^IgD^-^GL7^+^CD38^l^°) specific for spike (S^+^RBD^-^) or RBD (S^+^RBD^+^) probes. **(F)** Frequencies of TFH cells (CXCR5^++^BCL-6^+^CD4^+^CD3^+^B220). P values were derived by Mann-Whitney U tests. Data is presented as median ± IQR.

The profile of B and T cell immunity was assessed in draining lymph nodes 2 weeks after the boost immunisation. R-R elicited the lowest GC B cell responses measured using BCL-6^+^ (0.5%; IQR 0.4-1.2) or surface GL7^+^ (22.3%; IQR 20.5-31.6) expression (Figure 2D). Both S-S (BCL-6^+^: 1.9%; IQR 1.5-2.5) (GL7^+^: 47.8%, IQR 44.3-62.3) and S-R (BCL-6^+^: 1.9%; IQR 1.4-2.2) (GL7^+^: 52.7%; IQR 49.7-54.5) displayed intermediate induction of GC activity, while R-S immunised animals showed robust GC induction (BCL-6^+^: 3.0%; IQR 2.3-3.2) (GL7^+^: 54.2%; IQR 45.9-57.6), likely reflecting a primary response against non-RBD epitopes within S. The hierarchy of GC activity was mirrored in the frequency of probe-specific GC B cells, with R-S (19.5% (IQR:14.7-21.8) and S-S (10.6%; IQR 9.46-14.4) eliciting high frequencies of S-specific GC B cells (Figure 2E), while low frequencies were observed for R-R (1.83%; IQR 1.25-2.98) and S-R groups (2.9%; IQR 2.21-4.96). RBD-specific GC B cells were highest in R-R (8.07%; IQR 3.58-11.9) and S-R (6.79%; IQR 5.65-14.2) groups, with lower frequencies after S-S (2.86%; 0.96-7.11%) and R-S regimens (1.20%; IQR 0.67-2.62). GC TFH frequencies were highest in R-S animals (0.69%, IQR 0.6-1.1) compared to the other groups (R-R 0.4%, IQR 0.2-0.6; S-S 0.4%, IQR 0.3-0.4%; S-R 0.5%, IQR 0.2-0.7) (Figure 2F). The distribution of GC B and TFH responses after prime-boost immunisation were consistent in BALB/c mice (Figure S6).

### Both RBD and S immunogens elicit potent antibody responses and serum neutralising activity in non-human primates

To enable serial measurements in a highly relevant animal model, we next immunised pig-tailed macaques (*Macaca nemestrina*) with S and RBD protein vaccines formulated with an MPLA liposomal adjuvant (Baldrick et al., 2002). Two doses in all three vaccine regimens, R-R (n = 2), S-S (n = 3) and S-R (n = 3), reliably elicited robust serum antibody responses against S and RBD proteins (Figure 3A), together with a corresponding rise in both ACE2-RBD inhibition (Figure 3B) and neutralising activity (Figure 3C; median titre 202, IQR 160-241). Responses in macaques were notably more variable than mice and no major differences were discernible between the small groups.

**Figure 3.**
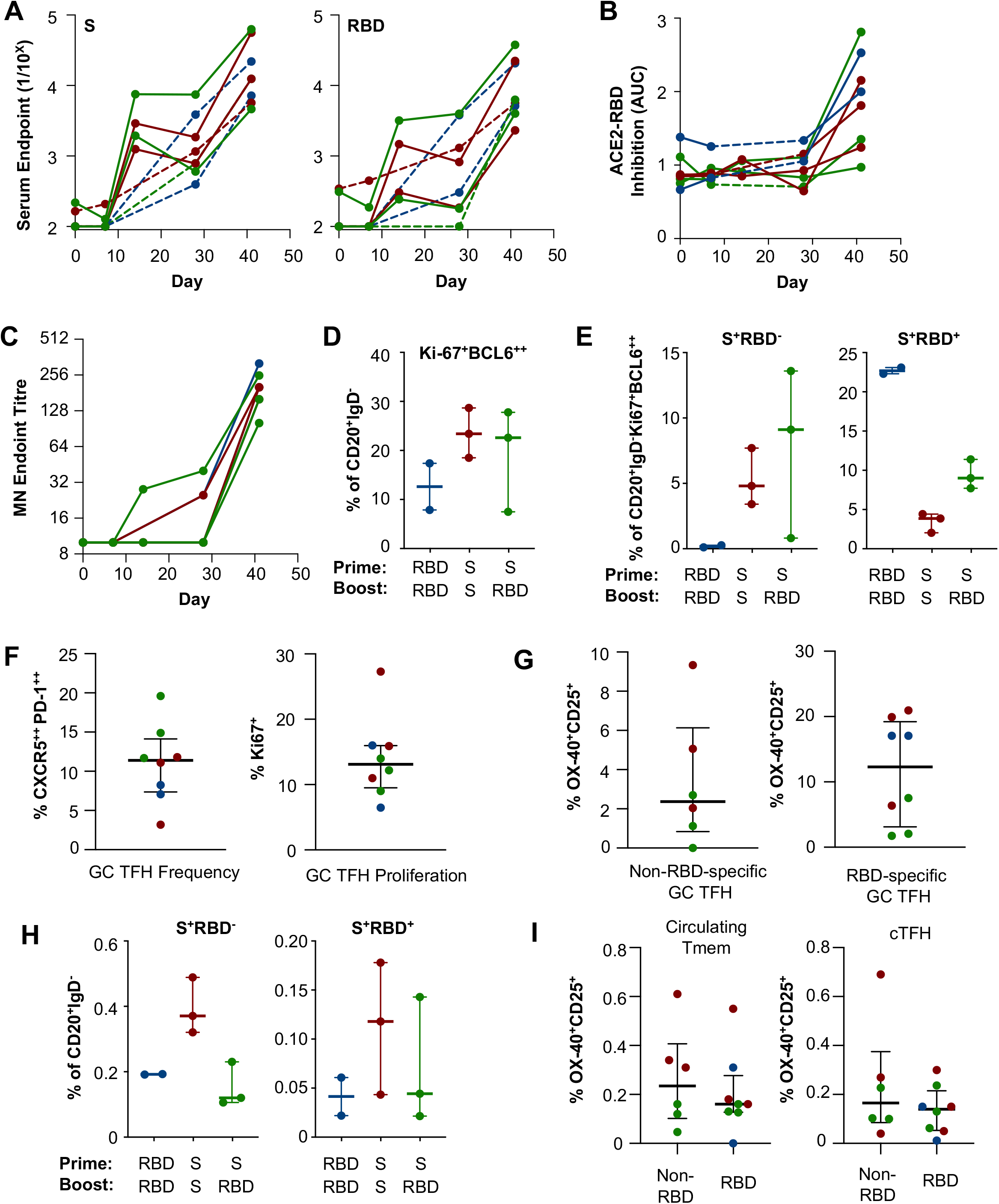
SARS-CoV-2 Spike and RBD immunogens elicit robust B and TFH responses in macaques. Macaques were serially immunised intramuscularly at a 28-day interval with S or RBD immunogens and immune responses assessed 13-14 days post-boost (R-R in blue (n=2), S-S in red (n=3) and S-R in green (n=3)). Dashed lines indicate animals without day 14 post-prime sampling. **(A)** Reciprocal endpoint titres of S- or RBD-specific IgG were measured in longitudinal plasma samples by ELISA. **(B)** The capacity of plasma antibodies to inhibit the interaction of RBD and human ACE2 was assessed by ELISA **(C)** Neutralisation activity in plasma was assessed using a microneutralisation assay. **(D)** Germinal centre CD20^+^IgD B cells were quantified based on Ki-67^+^BCL-6^+^ expression and frequencies of **(E)** spike-(S^+^RBD) or RBD-specific (S^+^RBD^+^) populations determined by flow cytometry. **(F)** Frequency of germinal centre TFH cells (CD3^+^CD4^+^CXCR5^++^PD-1^++^) and Ki-67^+^ expression in draining lymph nodes. **(G)** Antigen-specific TFH cells identified by OX-40^+^CD25^+^ upregulation following 18 hours of stimulation with S (minus RBD) or RBD peptide pools. **(H)** Frequency of circulating memory B cells within PBMC specific for spike (S^+^RBD) or RBD (S^+^RBD^+^). **(I)** Antigen-specific circulating memory CD4 T cells (Tmem; CD3^+^CD4^+^CD95^+^CXCR5^-^) or circulating TFH cells (cTFH; CD3^+^CD4^+^CD95^+^CXCR5^+^) identified as OX-40^+^CD25^+^ following 18 hours of stimulation with S (minus RBD) or RBD peptide pools. Antigen-specific T cell responses **(G and I)** are presented after background subtraction using a DMSO control. Data is presented as median ± IQR.

B and T cells responses to immunisation were assessed two weeks after the boost in the draining lymph nodes of immunised animals, identified based upon staining with a co-formulated tracking dye (Barber-Axthelm et al., 2020). The utility of employing a tracking dye for identifying vaccine-draining LN was exemplified by animal NM224 (Figure S7). While 7/8 animals exhibited dye staining of the iliac LN, the primary site of antigen drainage from quadricep intramuscular vaccination (Barber-Axthelm et al., 2020), NM224 showed dye staining of only the inguinal LN. Total GC B and TFH frequencies, as well as antigen-specific GC B and TFH frequencies, were enriched in the dyed inguinal LN compared to the unstained iliac LN (Barber-Axthelm et al., 2020).

Robust GC B cell responses (Figure 3D) (CD20^+^IgD^-^Ki-67^+^BCL-6^+^; gating Figure S8A) were elicited in all animals, with S-S (23.4%, range 18.5-28.7) and S-R (22.6%, range 7.5-27.8) immunised animals displaying higher GC frequencies relative to R-R (12.6%, range 7.86-17.4) immunised animals. GC B cell specificity was also assessed by S- or RBD-probe binding (Figure 3E). RBD-specific B cells were detected in all groups, with R-R immunised animals displaying the highest levels (22.7%, range 22.3-23.1) followed by the S-R (9.02%, range 7.73-11.4) and S-S (3.88%, range 2.03-4.43) groups. In contrast, high frequencies of S-specific GC B cells were elicited in S-S (4.81%, range 3.41-7.71) and S-R (9.11%, range 0.82-13.6) immunised, but not R-R immunised animals.

GC TFH (CD3^+^CD4^+^CXCR5^++^PD-1^++^; gating in Figure S9A) were also detected in all draining lymph nodes (11.4% of total LN CD4^+^ T cells), with a median of 13% exhibiting recent proliferation as measured by Ki-67 expression (Figure 3F). S-specific TFH targeting either non-RBD or RBD peptides were detected in all animals; interestingly, RBD-derived peptides tended to be more frequently recognised by GC TFH than non-RBD epitopes (2.4% non-RBD, S-specific versus 6.2% RBD-specific; Figure 3G).

The elicitation of memory lymphocyte populations is a key aim for protective vaccines. Circulating S- and RBD-specific memory B cells (CD20^+^IgD^-^; gating Figure S9B) were assessed in PBMCs (Figure 3H). S-specific memory B cells were highest in S-S immunised animals (0.37%, range 0.32-0.49), and approximately equivalent frequencies in R-R (0.195%, range 0.192-0.193), and S-R (0.12%, range 0.11-0.23) immunised animals. In contrast, RBD-specific memory B cells were less frequently detected overall, with S-S (0.12%, range 0.04-0.18) animals displaying the highest level, followed by the S-R (0.04%, range 0.02-0.14) and R-R (0.04%, range 0.02-0.06) groups. Both S-specific memory CD4 T cells (Tmem) and circulating TFH (cTFH; gating Figure S9B-C) were detected in blood two weeks after the vaccine boost (Figure 3I). In contrast to previous observations in convalescent human donors (Juno), NHP S-specific cTFH recognised non-RBD and RBD-derived peptides at similar frequencies (Figure 3J). Among all animals, RBD-specific cTFH frequencies correlated with S antibody titres suggesting that, analagous to humans (Bentebibel et al., 2013, 2016; Farooq et al., 2016; Huber et al., 2020; Koutsakos et al., 2018), antigen-specific cTFH constitute a useful biomarker of vaccine immunogenicity in NHP models (p=0.04, Figure S9D).

### Profile of responses in mice, macaques and humans

While our results suggest that adjuvanted protein vaccines are reliably immunogenic in mice and non-human primates, it remains unclear how well these models recapitulate human immunity to SARS-CoV-2. As a reference, S- and RBD-specific antibody and serum neutralisation titres were assessed in a panel of 72 convalescent donors recovered from COVID-19 (participant details in Table 1). We find that both S-S and S-R immunised mice display high comparative titres, with potent serum neutralisation appearing upon boosting with SARS-CoV-2 immunogens (Figure 4). Similarly, immunisation of NHP with one or two doses of SARS-CoV-2 immunogens, elicited binding antibodies at levels above the median observed in convalescent individuals. Neutralising responses in NHPs exceeded levels observed in convalescent individuals upon boosting with SARS-CoV-2 immunogens.

**Figure 4.**
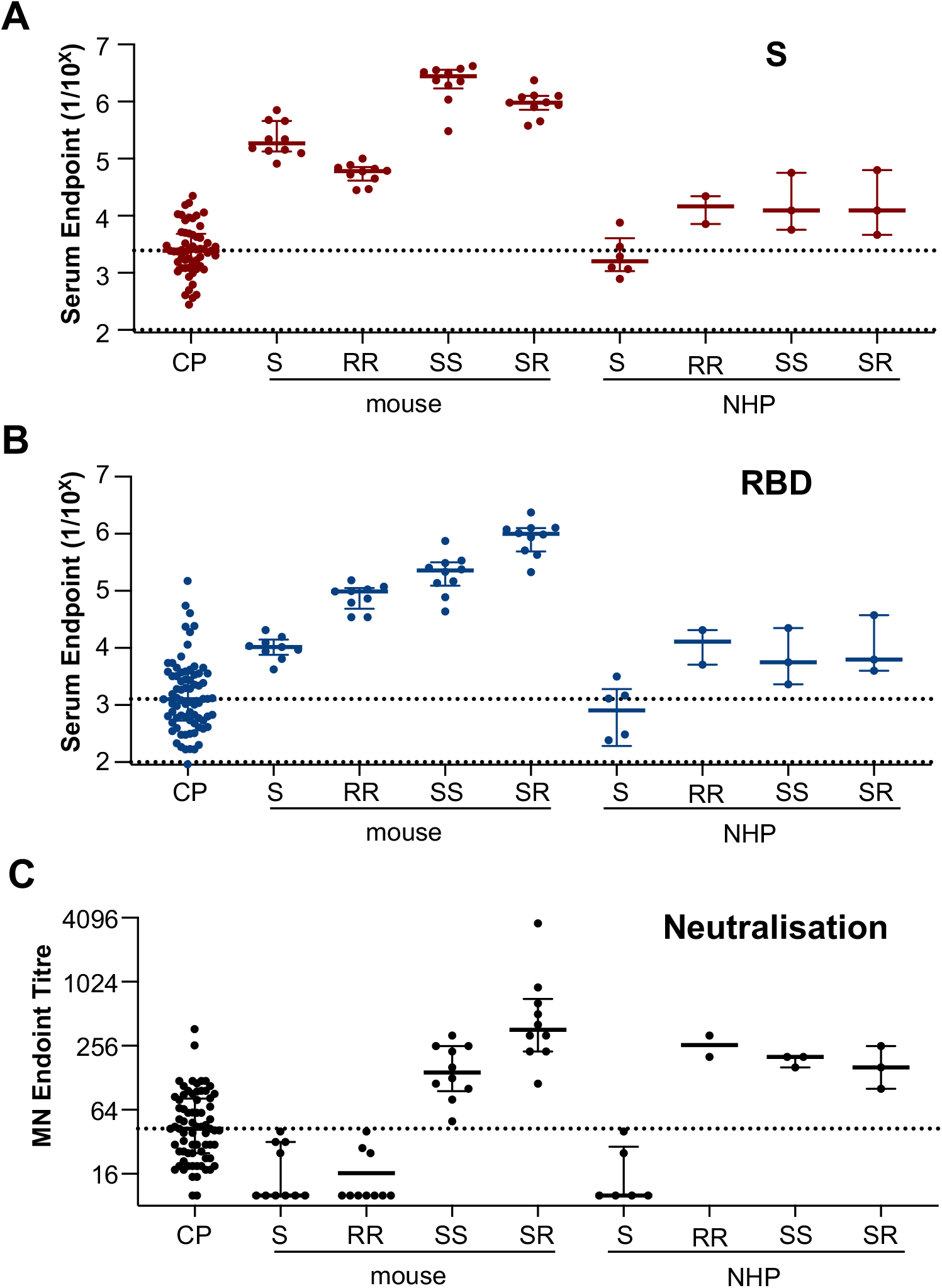
Comparative serological antibody and neutralising activity against SARS-CoV-2 across mice, macaques and convalescent humans. Serum from mice (n = 10 per group), and plasma from macaques (n = 2-3 per group) and human convalescent donors (n = 73) were assessed for endpoint total IgG titres measured by ELISA against **(A)** S or **(B)** RBD. **(C)** Neutralisation activity in the plasma (human, macaques) or serum (mouse) was assessed using a microneutralisation assay. **(A, B, C)** Dotted lines represent median of human convalescent donors.

**Table 1.** Characteristics of COVID-19 convalescent cohort

|  | Convalescent Subjects<br>(n=72) |
| --- | --- |
| Proportion female, % (n) | 41.6% (30) |
| Age, median (IQR) | 56 (62, 49) |
| Time since positive PCR test, median (IQR)* | 36 (31.5, 45) |
| Illness severity^, % (n) - mild | 68% (49) |
| - moderate | 22% (16) |
| - severe | 8% (6) |

\*Negative PCR test result for 6 individuals, all subjects seropositive
^Available for 71/72 participants

To examine the influence of host immunogenetics on antibody and B cell responses, we sorted and sequenced S- and/or RBD-specific B cells from convalescent subjects and immunised macaques, and germinal centre B cells from immunised mice. We find vaccine-elicited GC B cells in mice are drawn primarily from VH1-like gene families, in comparison to the VH3 and VH4 families that are predominant in RBD- and S- specific B cells recovered in the LN or PBMC of an immunised macaque (Figure S10). In contrast to immunised animals, RBD- and S-specific B cells sequenced from PBMC of convalescent subjects are drawn from a range of V-gene families, although we and others have reported biasing toward some VH3 family genes including VH3-30 and VH3-53/VH-3-66 (Juno et al., 2020; Robbiani et al., 2020). In terms of CDR-H3 length, recovered CDR-H3 from immunised mice were consistently shorter (median 11; IQR 10-12) relative to macaque LN (15; IQR 13-18) and PBMC (15; IQR 13-18), which were directly comparable to lengths observed for RBD- and S-specific immunoglobulin sequences observed in convalescent subjects (15; IQR 13-18).

## Discussion

Preliminary reports from SARS-CoV-2 vaccine candidates suggest both S (Folegatti et al., 2020; Jackson et al., 2020; Keech et al., 2020; Zhu et al., 2020) and RBD (Mulligan et al., 2020; Walsh et al., 2020) are immunogenic in human subjects. However the comparative performance of each immunogen in pre-clinical animal models, and the potential for combinatorial use in heterologous prime-boost strategies, is unclear. In line with other reports (Mandolesi et al., 2020; Walsh et al., 2020), we find the intrinsic immunogenicity of the RBD is limited in mice after a single or two doses, likely reflecting inefficient recruitment of high-quality TFH in the primary response, analogous to our previous report for influenza hemagglutinin stem immunogens (Tan et al., 2019). In contrast, priming of mice with S resulted in consistently higher serum titres and neutralising activity, irrespective of the subsequent boosting immunogen, highlighting the impact of a broader TFH repertoire in modulating the magnitude of immune responses after boosting. RBD was notably more immunogenic in vaccinated non-human primates in the context of greater MHCII loci and allelic diversity compared to mice. Nevertheless, the limited recognition of the RBD by B and cTFH cells following SARS-CoV-2 infection (Juno et al., 2020) suggests that at a population level, RBD-based immunogens might be less reliable vaccine antigens than S, which was robustly immunogenic in all species.

Selective recall of RBD responses by heterologous S-R prime boost immunisation focussed antibody recognition onto the ACE2 recognition site in mice, as evidenced by enhanced ACE2-RBD inhibition activity and a minor increase in neutralising activity. In contrast, enhanced neutralisation activity was not seen in S-R immunised macaques, where titres were equivalent to S-S animals. This mirrors our recent findings in convalescent humans, where serological neutralising activity was not exclusively RBD-directed (Juno et al., 2020), and suggests alternative S-localised epitopes such as the NTD (Chi et al., 2020), may contribute to vaccine-elicited protection. While mice have tremendous utility for immunological research including vaccine development efforts, we found divergent gene family usage and constrained CDR-H3 lengths in vaccine-elicited B cell responses in mice compared to immunised macaques or convalescent humans. These differences might explain the comparatively high serological binding titres required in mice for neutralisation activity compared to humans and immunised primates. Notably, many reported neutralising human monoclonal antibodies specific for epitopes outside the RBD, such as the N-terminal domain (Chi et al., 2020; Liu et al., 2020), display extended CDR-H3 loops of 20-25AA, considerably longer than what is commonly seen in mice. Therefore, genetic constraints in the murine immunoglobulin repertoire might impact vaccine-elicited humoral immunity to S, with implications for stratification of SARS-CoV-2 vaccines for eventual human use.

Most reports of clinical and pre-clinical SARS-CoV-2 vaccine assessment have been benchmarked against convalescent subjects recovered from COVID-19. However, numerous groups including ours have reported that neutralising titres in convalescent subjects are comparatively weak (Jackson et al., 2020; Juno et al., 2020; Keech et al., 2020; Mulligan et al., 2020; Walsh et al., 2020) and appear to wane rapidly (Ibarrondo et al., 2020; Long et al., 2020; Seow et al., 2020). Here a simple two protein regime using licensure-friendly adjuvants was able to elicit superior binding and neutralising antibody responses. Prototypic vaccines induced strong GC activity in draining lymph nodes, driving somatic maturation of S-specific B cells, and seeded memory T and B cell responses in the blood. Overall, our study suggests that vaccination constitutes a more robust and reliable pathway to serological protection against SARS-CoV-2 than natural infection, similar to other pathogens such as human papillomavirus (Harper et al., 2004; Villa et al., 2006).

## Acknowledgements

We thank the generous participation of the trial subjects for providing samples. The SARS-CoV-2 RBD expression plasmids were kindly provided by Florian Krammer, Mt Sinai School of Medicine, NY, USA. The human and mouse ACE2 expression plasmids were kindly provided by Merlin Thomas, Monash University, Australia. We acknowledge the Melbourne Cytometry Platform (Melbourne Brain Centre node) for provision of flow cytometry services. We thank Robin Shattock (Imperial College London) and Dietmar Katinger and Philipp Mundsperger from Polymun for provision of the MPLA liposome adjuvant. This study was supported by the Victorian Government and Medical Research Future Fund (MRFF) GNT2002073 (SJK and AKW), the ARC Centre of Excellence in Convergent Bio-Nano Science and Technology (SJK), an NHMRC program grant APP1149990 (SJK), NHMRC project grant GNT1162760 (AKW), an NHMRC-EU collaborative award APP1115828 (SJK), the European Union Horizon 2020 Research and Innovation Programme under grant agreement 681137 (SJK), the Jack Ma Foundation (KS) and the A2 Milk Company (KS). JAJ and SJK are supported by NHMRC fellowships. AKW and KS are supported by NHMRC Investigator grants. The Melbourne WHO Collaborating Centre for Reference and Research on Influenza is supported by the Australian Government Department of Health.

## Author Contributions

HXT, JAJ, WSL, SJK and AKW designed the study and experiments; HXT. JAJ, WSL, IBA, HGK, KW, RE, TA, FLM and AKW performed experiments; KS contributed unique reagents; HXT, JAJ, WSL, KS, SJK and AKW analysed the experimental data; HXT, JAJ, WSL, SJK and AKW wrote the manuscript. All authors reviewed the manuscript.

## Materials and Methods

### Ethics Statement

Animal studies and related experimental procedures were approved by the University of Melbourne Animal Ethics Committee (#1714193, #1914874). Macaque studies and related experimental procedures were approved by the Monash University Animal Ethics Committee (#23997). Human clinical study protocols were approved by the University of Melbourne Human Research Ethics Committee (#2056689), and all associated procedures were carried out in accordance with approved guidelines. All participants provided written informed consent in accordance with the Declaration of Helsinki.

### Expression of coronavirus proteins

The recombinant expression and validation of SARS-CoV-2 spike and RBD proteins was previously described (Juno et al., 2020). Briefly, the ectodomain of SARS-CoV-2 (isolate WHU1; residues 1 – 1208) with furin cleavage site removed and P986/987 stabilisation mutations^36^, a C-terminal T4 trimerisation domain, Avitag and His-tag, was expressed in Expi293 cells and purified by Ni-NTA size-exclusion chromatography. The SARS-CoV-2 RBD^38^ with a C-terminal His-tag (residues 319-541; kindly provided by Florian Krammer) was expressed in Expi293 cells and purified by Ni-NTA.

### Animal immunisations

5μg of S, RBD or OVA proteins were formulated in PBS at a 1:1 ratio with Addavax adjuvant (InvivoGen). C57BL/6 or BALB/c mice were anesthetised by isoflurane inhalation prior to intramuscular injection of 50 μL vaccine in each hind quadriceps. Primary responses were assessed 14 days after prime immunisation. Booster immunisations were administered 3 weeks post-prime, and responses assessed 14 days after boost.

Pigtail macaques (*Macaca* nemistrina) were housed in the Monash Animal Research Platform and animals were recycled from a preceding gamma delta (γδ) T cell immunotherapy trial after confirmation γδ T cell frequencies had returned to baseline levels. Eight male macaques (*Macaca nemestrina*) (6-15 years old) were vaccinated with 100μg of SARS-CoV-2 spike or RBD immunogens formulated with 200μg of Monophosphoryl Lipid A (MPLA) liposomes (Polymun) (Baldrick et al., 2002) intramuscularly in the right quadriceps. 28 days after priming, booster immunisations consisting of 100μg S or RBD protein with 200μg of MPLA and 1% tattoo ink were administered intramuscularly in both quadriceps. Although unlikely to be confounding, macaques were concurrently vaccinated in the right and left deltoids with HIV-trimeric envelope protein gp140 (SOSIP) immunogens (100μg) (Ringe et al., 2013; Sliepen et al., 2019) formulated with MPLA and 1.0% tattoo ink (right deltoid only). Twenty-four hours prior to necropsy, macaques received an intravenous infusion of autologous Vδ2^+^Vγ9^+^ T-cells labelled with CellTrace Blue (Life Technologies). Any CellTrace Blue^+^ cells were excluded from flow cytometric analysis of B or T cell populations.

### Flow cytometric detection of S and RBD-specific B cells

S protein was biotinylated using Bir-A (Avidity) and labelled by the sequential addition of streptavidin (SA) conjugated to PE (BD). RBD protein was directly labelled to APC using an APC Conjugation Lightning-Link Kit (Abcam). For murine studies, lymph nodes were mechanically homogenised into single cell suspensions in RF10 media (RPMI 1640, 10% FCS, 1× penicillin-streptomycin-glutamine; Life Technologies). Isolated cells were stained with Aqua viability dye (Thermofisher) and Fc-blocked with a CD16/32 antibody (93; Biolegend). Cells were then surface stained with S/RBD probes and the following antibodies: B220 BUV737 (RA3-6B2), IgD BUV395 (11-26c.2a), CD45 Cy7APC (30-F11), SA BV786 (BD), GL7 Alexa488 (GL7), CD38 Cy7PE (90), CD3 BV786 (145-2C11) and F4/80 BV786 (BM) (Biolegend). Cells were washed twice with PBS containing 1% FCS and fixed with 1% formaldehyde (Polysciences).

For macaque studies, cryopreserved single cell suspensions were thawed, and stained with Aqua viability dye (Thermofisher). Cells were then surface stained with S/RBD probes and the following antibodies: IgD AF488 (polyclonal; Southern Biotech), IgM BUV395 (G20-127), IgG BV786 (G18-145) (BD), CD14 BV510 (M5E2), CD3 BV510 (OKT3), CD8a BV510 (RPA-T8), CD16 BV510 (3G8), CD10 BV510 (HI10a), CD20 APC-Cy7 (2H7) (Biolegend) and SA BV510 (BD). Cells were washed twice with PBS containing 1% FCS and fixed with 1% formaldehyde (Polysciences). For intracellular transcription factor staining, cells were first stained with Aqua viability dye (Life Technologies), followed by S/RBD probes and surface antibodies: IgD AF488 (polyclonal; Southern Biotech), IgG BV786 (G18-145) (BD), CD14 BV510 (M5E2), CD3 BV510 (OKT3), CD8a BV510 (RPA-T8), CD16 BV510 (3G8), CD10 BV510 (HI10a), CD20 APC-Cy7 (2H7) (Biolegend) and SA BV510 (BD). Cells were washed and permeabilised with Transcription Factor Buffer Set (BD) prior to BCL-6 PE-Cy7 (K112-91, BD) and Ki-67 BUV395 (B56, BD) staining. Cells were washed twice and resuspended in PBS containing 1% FCS. Samples were acquired on a BD LSR Fortessa using BD FACS Diva.

### Flow cytometric detection of ex vivo and antigen-specific TFH

For *ex vivo* TFH quantification from mice, freshly isolated LN single cell suspensions were stained with the following antibodies: Live/dead Red (Life Technologies), CD3 BV510 (145-2C11; Biolegend), PD-1 BV786 (29F.1A12; Biolegend), CXCR5 BV421 (L138D7; Biolegend), CD4 BUV737 (RM4-5; BD), B220 BV605 (RA3-6B2; BD), and F4/80 PE-Dazzle 594 (T45-2342; BD). Cells were permeabilized with transcription factor staining buffer (BD Biosciences) and stained intracellularly with anti-BCL-6 Alexa647 (IG191E/A8; Biolegend). Non-human primate LN suspensions and PBMC were stained with the same protocol, using the following antibodies: Live/dead Aqua (Life Technologies), CD3 Alexa700 (SP34-2; BD), PD-1 BV421 (EH12.2H7; Biolegend), CXCR5 PE (MU5UBEE; ThermoFisher), CD4 BV605 (L200; BD), CD20 BV510 (2H7; BD), CD8 BV650 (RPA-T8; Biolegend), CD95 BUV737 (DX2; BD), ICOS PerCP-Cy5.5 (C398.4A; Biolegend), CD69 FITC (FN50; Biolegend), CCR6 BV785 (G034E3; Biolegend), CXCR3 Pe-Dazzle594 (G02H57; Biolegend), BCL-6 APC (IG191E/A8; Biolegend) and Ki67 BUV395 (B56; BD).

To identify antigen-specific TFH cells, LN cell or PBMC suspensions were cultured in RF10 media for 18 hours at 37°C. Cryopreserved NHP samples were rested for 4 hours at 37°C prior to stimulation, while murine samples were processed fresh. Samples were stimulated with a peptide pool (15mers overlapping by 11, 2μg/peptide/mL) comprising the SARS-CoV-2 RBD or SARS-CoV-2 S (without the RBD), or a DMSO control. At the time of stimulation, an anti-mouse CD154 PE mAb (MR1; Biolegend) or anti-human CD154 APC-Cy7 (TRAP1; BD) was added to all culture conditions. After stimulation, cells were washed twice in PBS and stained with viability dye (Red or Aqua, Life Technologies) according to the manufacturer’s instructions. Mouse cells were then stained with CD3 BV510 (145-2C11; Biolegend), CD25 BB515 (PD61; BD), PD-1 BV786 (29F.1A12; Biolegend), CXCR5 BV421 (L138D7; Biolegend), CD4 BUV737 (RM4-5; BD), OX-40 PeCy7 (OX-86; Biolegend), B220 BV605 (RA3-6B2; BD), and F4/80 PE-Dazzle 594 (T45-2342; BD) before being washed and fixed. NHP cells were stained with the following antibodies: CD3 Alexa700 (SP34-2; BD), PD-1 BV421 (EH12.2H7; Biolegend), CXCR5 PE (MU5UBEE; ThermoFisher), CD4 BV605 (L200; BD), CD20 BV510 (2H7; BD), CD8 BV650 (RPA-T8; Biolegend), CD95 BUV737 (DX2; BD), CCR6 BV785 (G034E3; Biolegend), CXCR3 Pe-Dazzle594 (G02H57; Biolegend), CD25 APC (BC96; Biolegend), and OX-40 BUV395 (L106; BD). Samples were acquired on a BD LSR Fortessa using BD FACS Diva.

### B cell receptor sequencing and analysis

B cell receptor sequences were recovered from GC B cells (B220+IgD-GL7+) in the draining iliac lymph node of C57BL/6 mice (n=3) 14 days after a single immunisation with S. Single cells were sorted using a BD Aria II into 96 well plates and subject to cDNA generation and multiplex PCR and sanger sequencing as previously described (Tiller et al., 2009). For macaques, a single cell suspension was prepared from the draining iliac lymph node of a single animal 14 days after a second immunisation with S. Single S-(S+RBD-) and RBD-specific (S+RBD+) IgG+ B cells were stained as above and sorted using a BD Aria II into 96 well plates, subject to cDNA generation and multiplex PCR and sanger sequencing using previously described techniques (Mason et al., 2016). Productive, recombined heavy (V-D-J) and light chain (V-J) immunoglobulin sequences were analysed using IMGT V-quest (Brochet et al., 2008).

### ELISA

Antibody binding to coronavirus S or RBD proteins was tested by ELISA. 96-well Maxisorp plates (Thermo Fisher) were coated overnight at 4°C with 2_μg/mL recombinant S, RBD or OVA proteins. After blocking with 1% FCS in PBS, duplicate wells of serially diluted plasma were added and incubated for two hours at room temperature. Plates were washed prior to incubation with HRP-conjugated secondary antibodies for mouse (1:10000; anti-mouse IgG; KPL), macaque (1:10000; anti-macaque IgG; Rockland) or human (1:20000; anti-human IgG; Sigma) for 1 hour at room temperature. Plates were washed and developed using TMB substrate (Sigma) and read at 450nm. Endpoint titres were calculated as the reciprocal serum dilution giving signal 2x background using a fitted curve (4 parameter log regression).

### ACE2-RBD inhibition ELISA

An ELISA to measure the ability of plasma antibodies to block interaction between recombinant human ACE2 (kindly provided by Merlin Thomas) and SARS-CoV-2 RBD was performed as previously described (Juno et al., 2020). Briefly, 96-well Maxisorp plates (Thermo Fisher) were coated overnight at 4°C with 2.5μg/ml of recombinant RBD protein in carbonate-bicarbonate coating buffer (Sigma). After blocking with PBS containing 1% BSA, duplicate wells of serially diluted plasma (1:25 to 1:102,400) were added and incubated for 1 hour at room temperature. Plates were then incubated with 1μg/ml of biotinylated recombinant ACE2 protein for 1 hour at room temperature followed by incubation with HRP-conjugated streptavidin (Thermo Fisher Scientific) for 1 hour at room temperature. Plates were developed with TMB substrate (Sigma), stopped with 0.15M sulphuric acid and read at 450nm. %Inhibition was plotted against plasma dilutions and the area under curve (AUC) was calculated using Graphpad Prism.

### Microneutralisation Assay

SARS-CoV-2 isolate CoV/Australia/VIC01/2020 (Caly et al., 2020) was passaged in Vero cells and stored at −80 °C. Plasma was heat inactivated at 56 °C for 30 min. Plasma was serially diluted 1:20 to 1:10,240 before the addition of 100 TCID_50_ of SARS-CoV-2 in MEM/0.5% BSA and incubation at room temperature for 1 h. Residual virus infectivity in the plasma/virus mixtures was assessed in quadruplicate wells of Vero cells incubated in serum-free media containing 1 μg/ml of TPCK trypsin at 37 °C and 5% CO2; viral cytopathic effect was read on day 5. The neutralising antibody titre was calculated using the Reed–Muench method, as previously described (Houser et al., 2016; Subbarao et al., 2004).

### Statistics

Data is generally presented as median +/- interquartile range or range. Statistical significance was assessed by Mann-Whitney U tests. Curve fitting was performed using 4 parameter logistic regression. All statistical analyses were performed using Prism (GraphPad). Flow cytometry data was analysed in FlowJo v9 or v10.

---

**Supplemental Figure 1.**
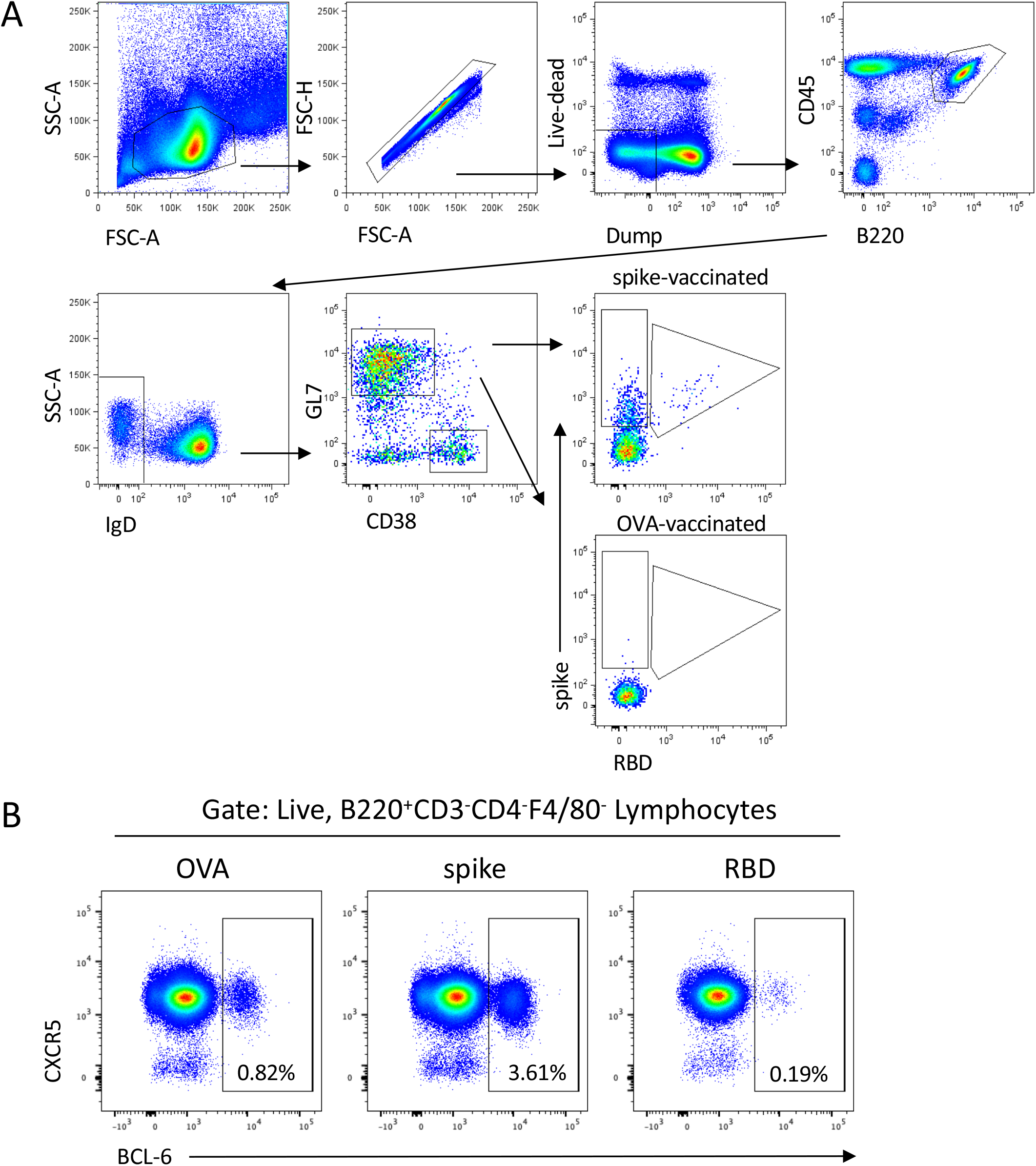
Identification of bulk and antigen-specific mouse germinal centre B cells. **(A)** Lymphocytes were identified by FSC-A vs SSC-A gating, followed by doublet exclusion (FSC-A vs FSC-H). Live and CD3-F4/80-streptavidin-(dump channel) cells were gated and CD45^+^B220^+^IgD^-^ B cells identified. Germinal centre (GL7^+^CD38^lo^) B cells were then assessed for binding to SARS-CoV-2 spike (S) and/or SARS-CoV-2 RBD probes. **(B)** Frequencies of germinal centre B cells was alternatively measured based on BCL-6 expression in live, B220^+^CD3^-^CD4^-^F4/80^-^ lymphocytes.

**Supplemental Figure 2.**
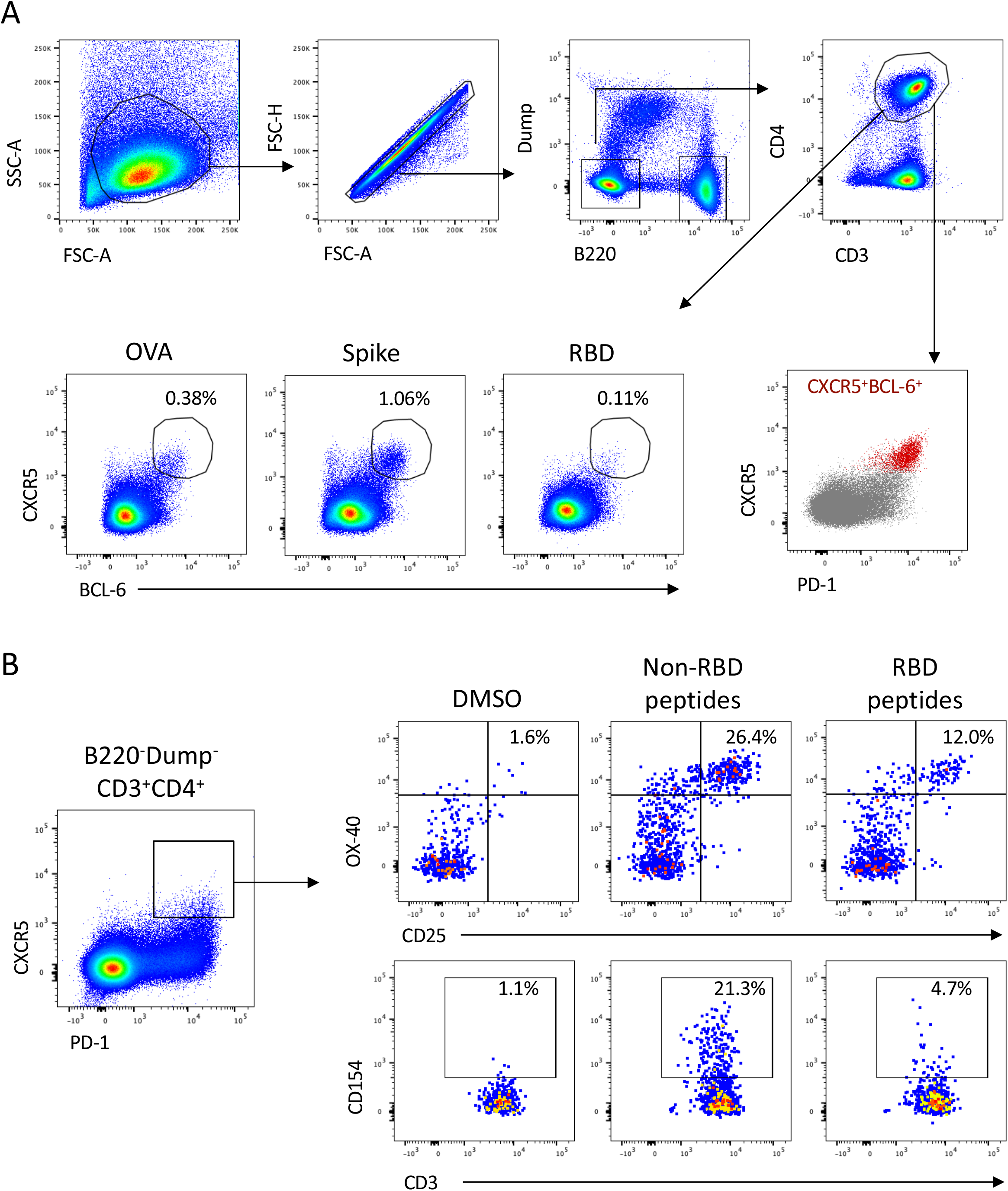
Identification of bulk and antigen-specific mouse TFH cells. **(A)** Gating strategy for ex vivo identification of lymph node TFH cells, based on BCL-6 and CXCR5 expression. Comparison of bulk CD4^+^ (grey) and TFH cells (red) confirms the PD-1^hi^ phenotype of the BCL-6^+^CXCR5^hi^ population. **(B)** Identification of S-specific TFH (OX-40^++^CD25^+^ or CD154^+^) following peptide pool stimulation of lymph node suspensions from S-vaccinated animals (plots represent pooled lymph nodes from 3 animals).

**Supplemental Figure 3.**
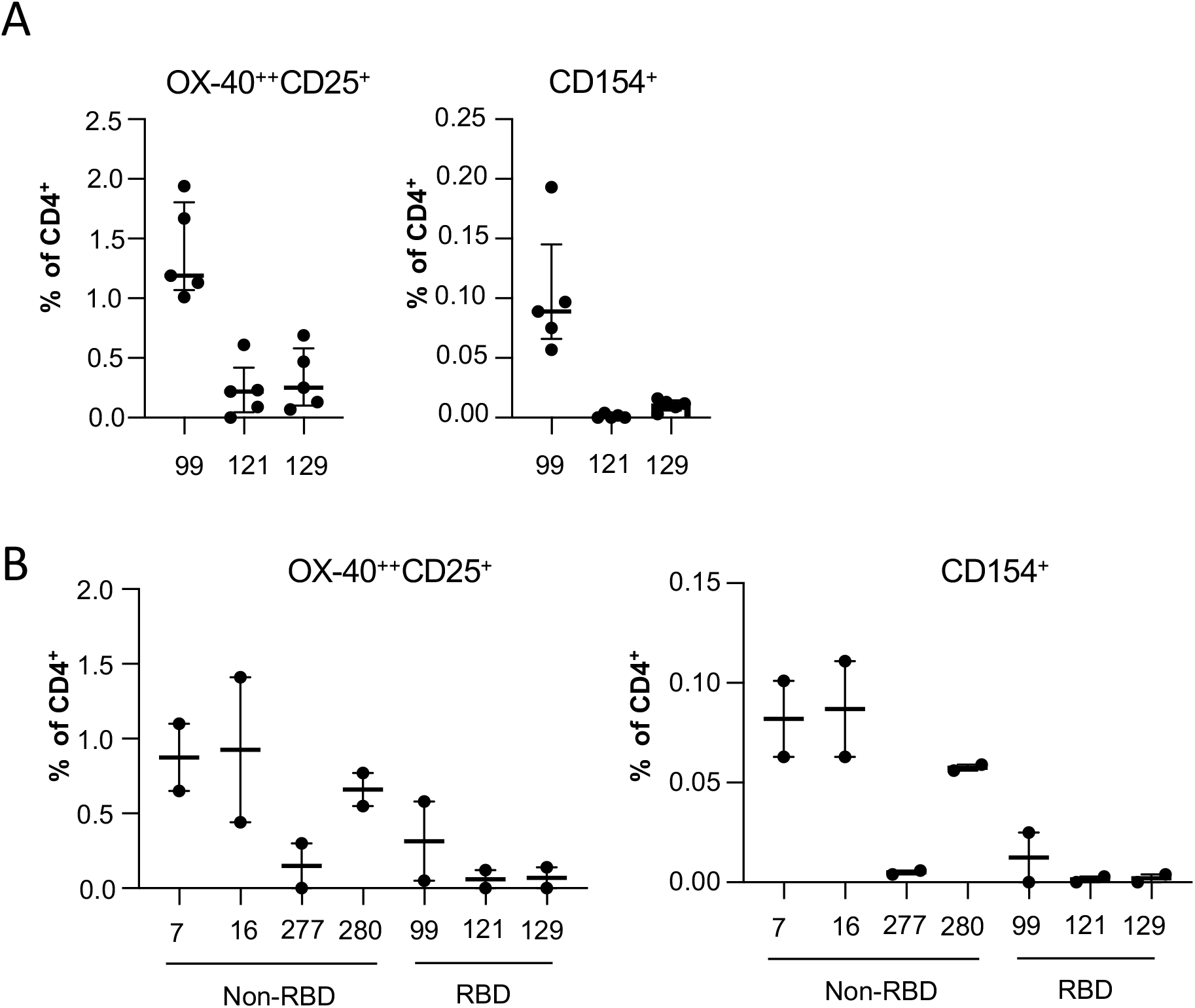
SARS-CoV-2 RBD immunogenic T cell epitopes. **(A)** Screening of RBD-derived 15-mer peptides identified 3 T cell epitopes recognized by C57BL/6 mice. Graphs indicate the frequency of OX-40^++^CD25^+^ or CD154^+^ CD4^+^ cells following in vitro peptide re-stimulation of lymph node suspensions from RBD-vaccinated mice (n=5 individual mice). **(B)** Screening of non-RBD peptides identified multiple T cell epitopes immunogenic in C57BL/6 mice, including peptides 7, 16, 277 and 280. Graphs indicate the frequency of peptide-specific CD4^+^ T cells in pooled lymph node suspensions of S-vaccinated mice (each data point represents a pool of 5 mice).

**Supplemental Figure 4.**
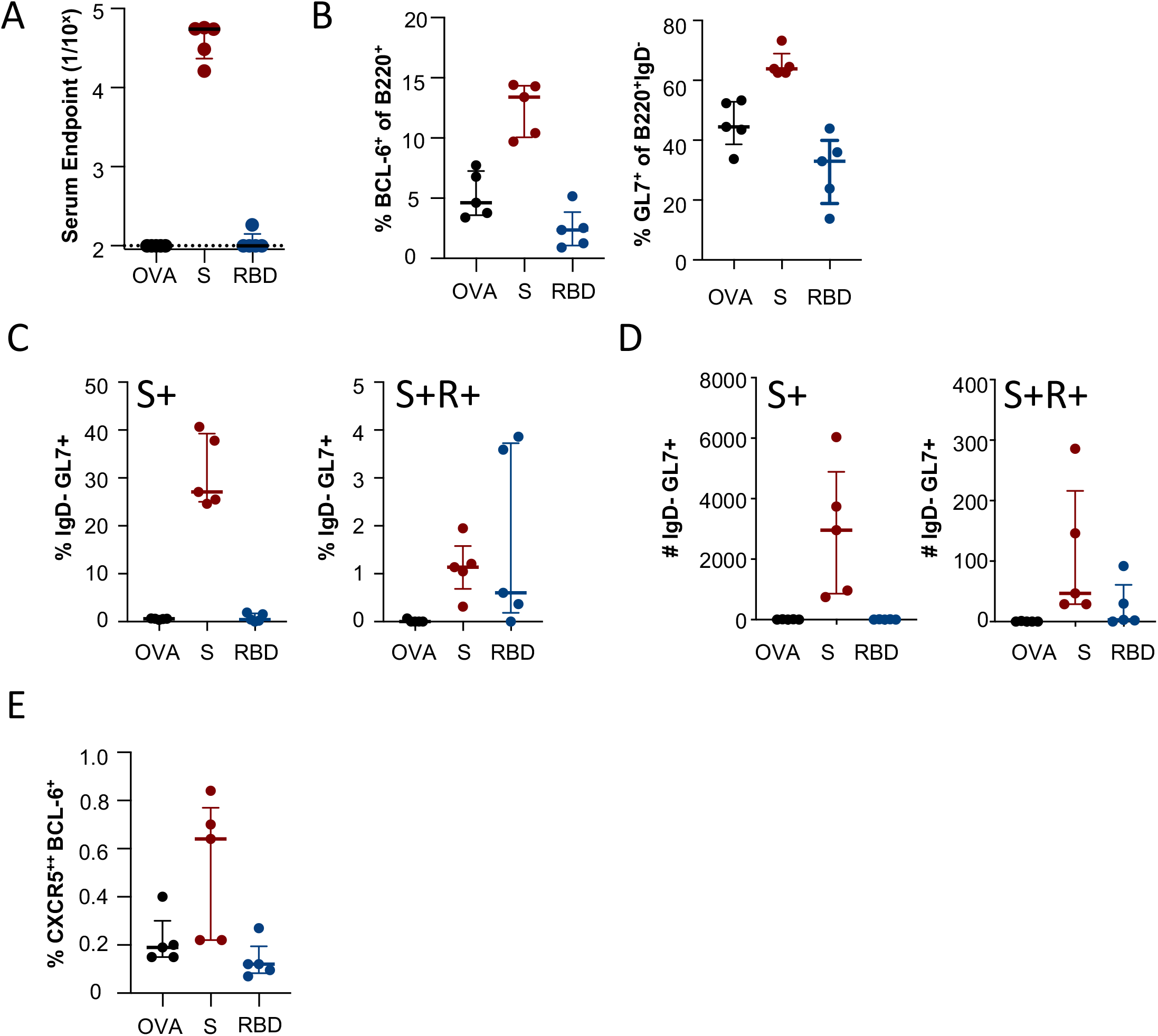
Primary immunogenicity of SARS-CoV-2 subunit proteins in BALB/c mice. Mice were immunised intramuscularly with S, RBD or OVA immunogens and immune responses assessed 14 days post-immunisation (n = 5). **(A)** Reciprocal serum endpoint dilutions of S-(red), RBD-(blue) or OVA-specific IgG (black) were measured by ELISA. Dotted lines denote the detection cut off (1:100 dilution). **(B)** Draining lymph node germinal centre activity assessed by BCL-6 expression in B220^+^ B cells or GL7 expression in B220^+^IgD^-^ B cells. **(C)** Frequency and **(D)** absolute counts of germinal centre B cells (B220^+^IgD^-^GL7^+^CD38^lo^) specific for spike (S^+^RBD^-^) or RBD (S^+^RBD^+^) probes. **(E)** Frequency of TFH cells (CXCR5^++^BCL-6^+^CD4^+^CD3^+^B220^-^). Data is presented as median ± IQR.

**Supplemental Figure 5.**
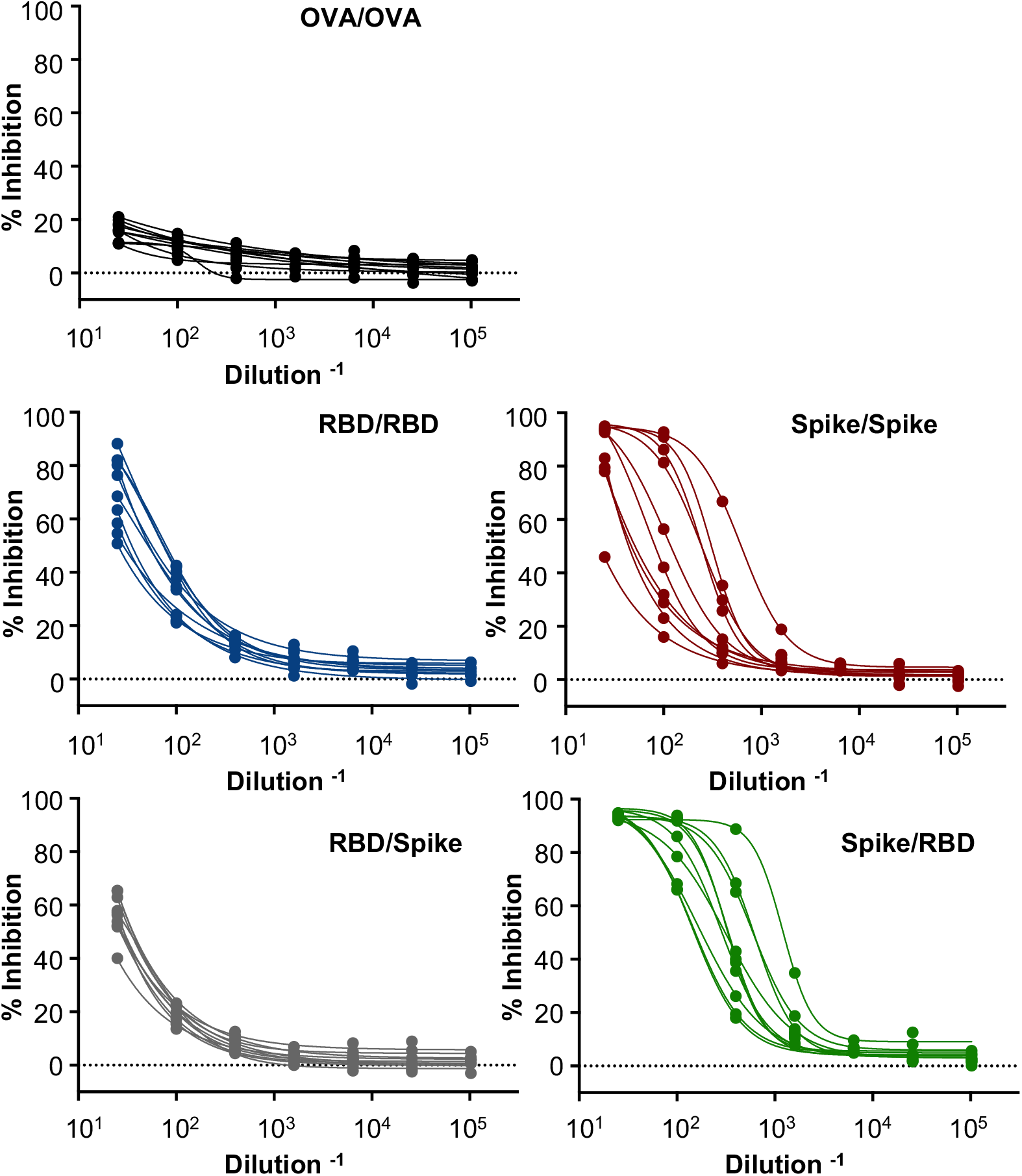
Inhibition of ACE2-RBD engagement by serum antibodies in immunised C57BL/6 mice. Mice were serially immunised intramuscularly at a 21-day interval with S, RBD or OVA proteins and immune responses assessed 14 days post-boost. The capacity for serum antibodies from immunised mice (n = 10 per group) to inhibit the interaction of recombinant SARS-CoV-2 RBD and human ACE2 was assessed by ELISA.

**Supplemental Figure 6.**
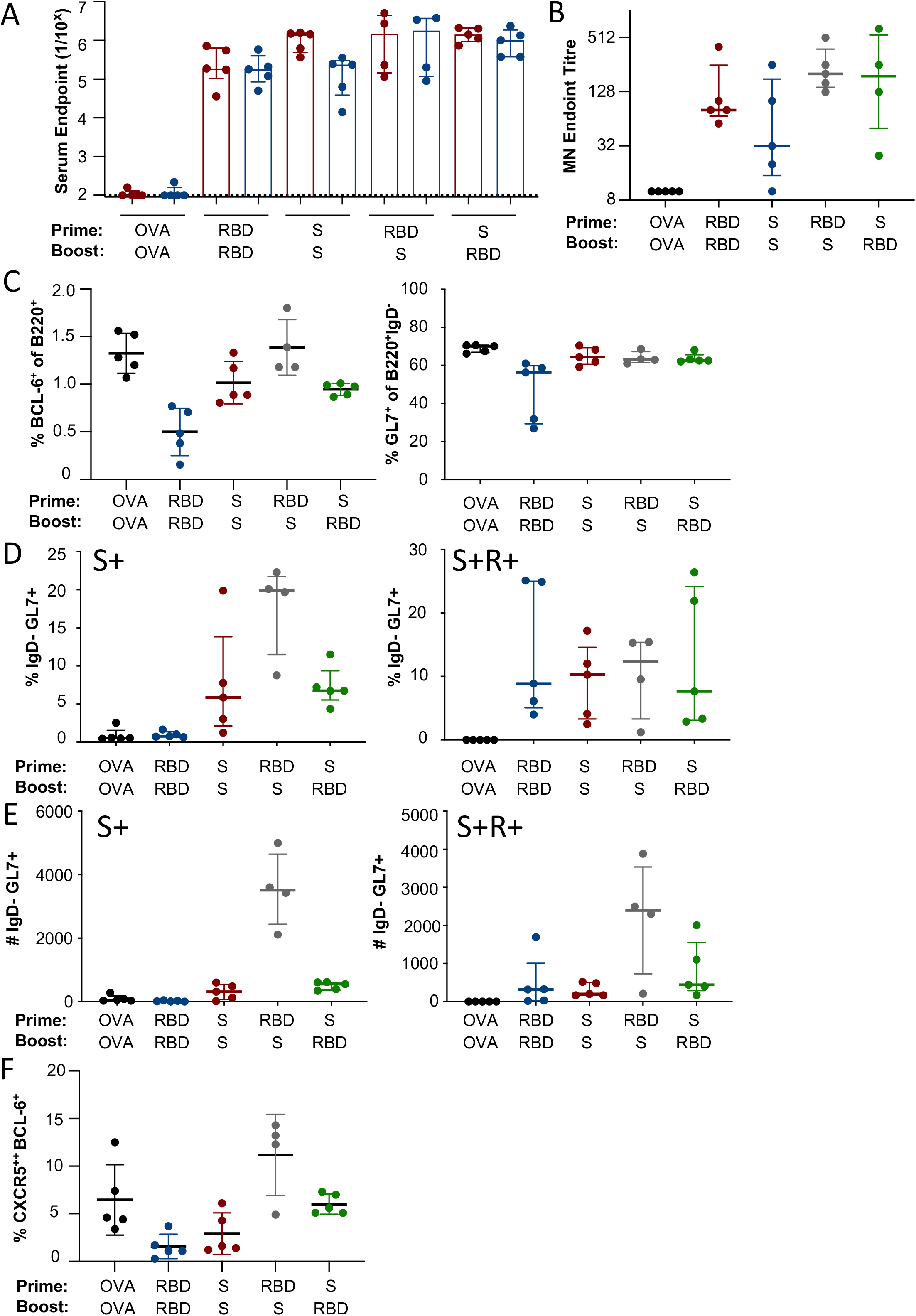
Prime-boost immunisation of SARS-CoV-2 subunit proteins in BALB/c mice. Mice were serially immunised intramuscularly at a 21-day interval with S, RBD or OVA proteins and immune responses assessed 14 days post-boost (n = 5). **(A)** Reciprocal serum endpoint dilutions of S-(red) or RBD-specific (blue) were measured by ELISA. Dotted lines denote the detection cut off (1:100 dilution). **(B)** Neutralisation activity in the serum was assessed using a microneutralisation assay. **(C)** Draining lymph node germinal centre activity assessed by BCL-6 expression in B220^+^ B cells or GL7 expression in B220^+^IgD^-^ B cells. **(D)** Frequency and **(E)** absolute counts of of germinal centre B cells (B220^+^IgD^-^GL7^+^CD38^lo^) specific for spike (S^+^RBD^-^) or RBD (S^+^RBD^+^) probes. **(F)** Frequency of TFH cells (CXCR5^++^BCL-6^+^CD4^+^CD3^+^B220^-^). Data is presented as median ± IQR.

**Supplemental Figure 7.**
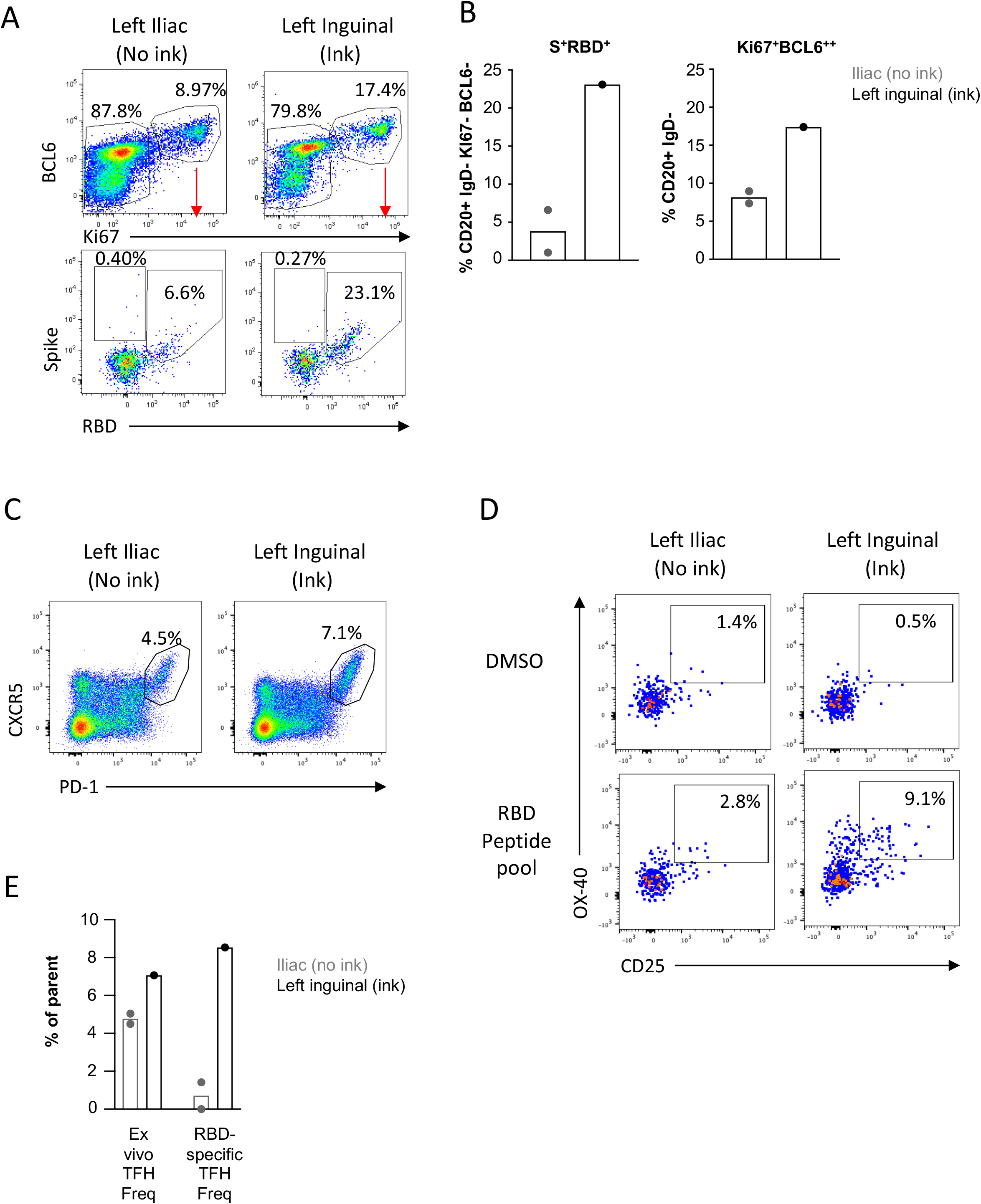
Enrichment of GC responses in ink-stained lymph nodes. **(A)** Ex vivo staining (gating in Figure S9A) and **(B)** frequency of RBD-specific germinal centre B cell in the left iliac or inguinal LN. **(C)** Ex vivo staining of GC TFH cells (gating in Figure S10A) in the left iliac or inguinal LN. **(D)** Quantification of RBD-specific TFH following peptide pool stimulation in left iliac or inguinal LN. **(E)** Summary of ex vivo bulk TFH frequency or RBD-specific TFH among left and right iliac LN (with no evidence of ink staining) or left inguinal LN (with ink staining).

**Supplemental Figure 8.**
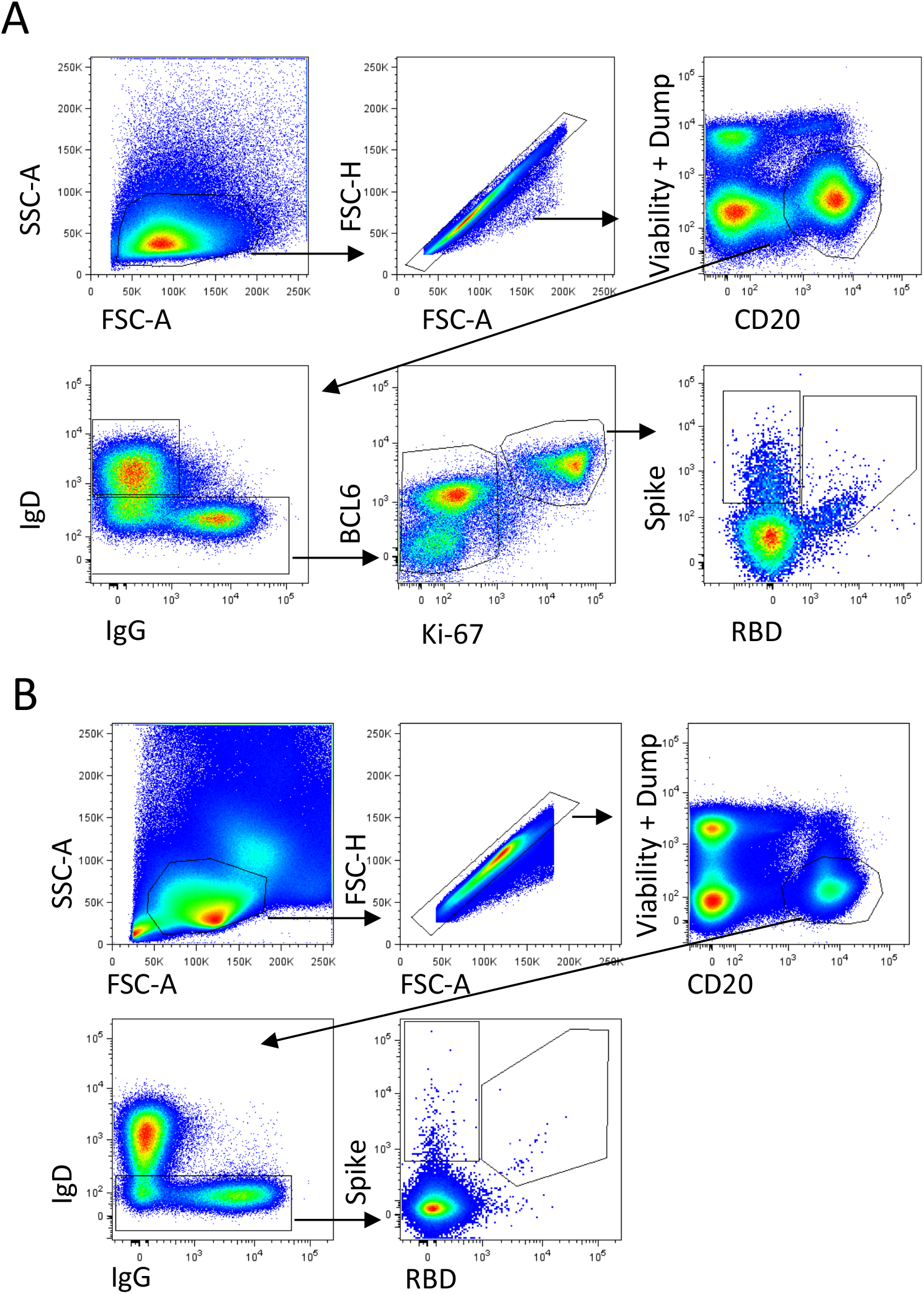
Macaque B cell gating strategy. Gating of antigen-specific **(A)** germinal centre B cells in lymph nodes or **(B)** circulating memory B cells in PBMC samples. Lymphocytes were identified by FSC-A vs SSC-A gating, followed by doublet exclusion (FSC-A vs FSC-H), and gating on dump^-^ (CD3^-^ CD8^-^CD14^-^CD10^-^CD16^-^streptavidin^-^) live CD20^+^ B cells. **(A)** Antigen-specific germinal center B cells were identified from class-switched IgD-B cells and intracellular expression of BCL6 and Ki-67. Alternatively, **(B)** circulating memory B cells in PBMC samples were identified as CD20^+^IgD^-^. Antigen specificity was determined by binding to SARS-CoV-2 spike (S) and/or SARS-CoV-2 RBD probes.

**Supplemental Figure 9.**
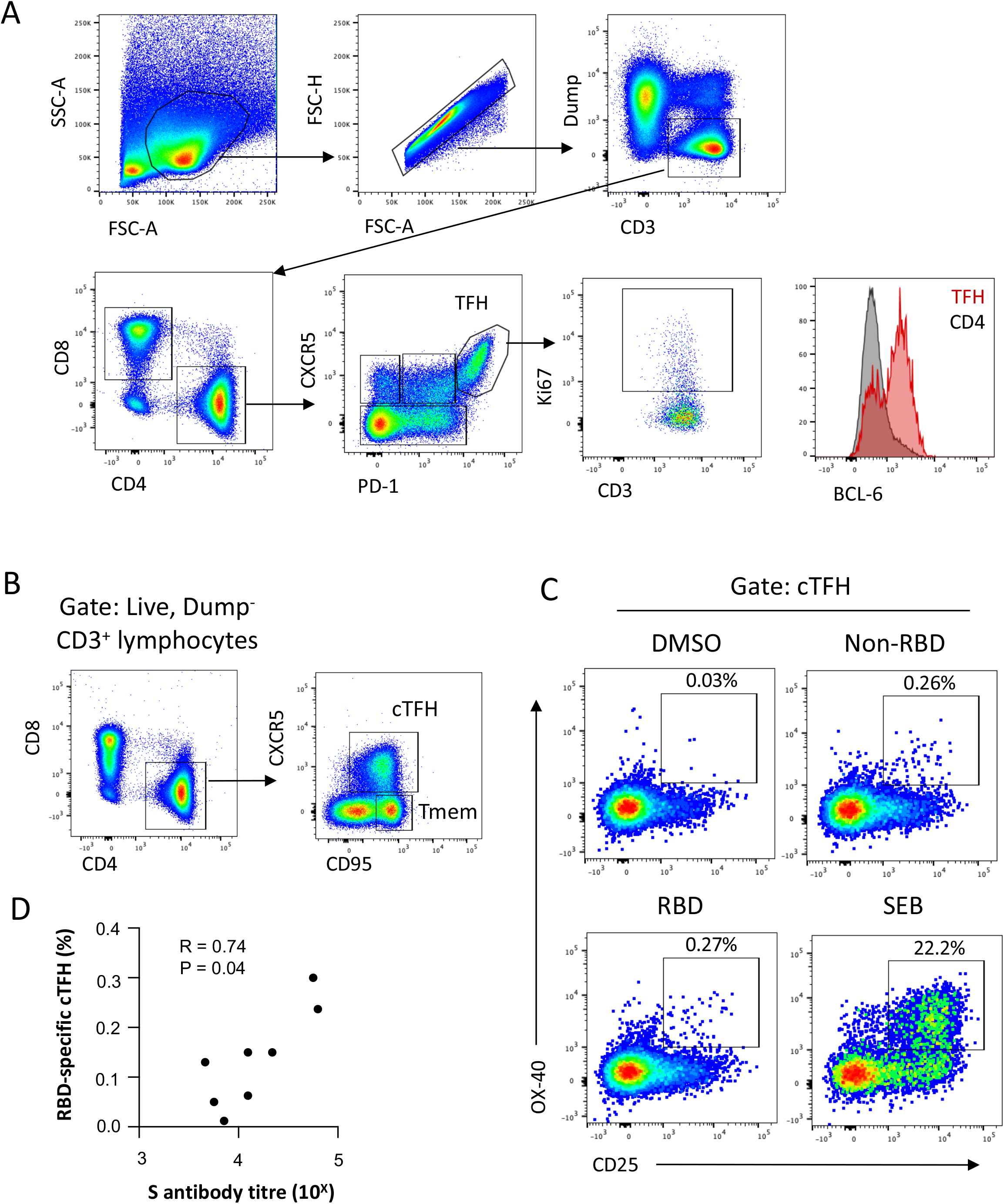
Macaque T cell gating strategy. **(A)** Gating of lymph node GC TFH (CXCR5^hi^PD-1^hi^) cells, and expression of Ki-67 and BCL-6. **(B)** Ex vivo identification of macaque cTFH in PBMC. **(C)** Identification of OX-40^+^CD25^+^ cTFH following in vitro peptide pool or SEB stimulation. **(D)** Spearman correlation of RBD-specific cTFH frequency and S antibody titres at D41.

**Supplemental Figure 10.**
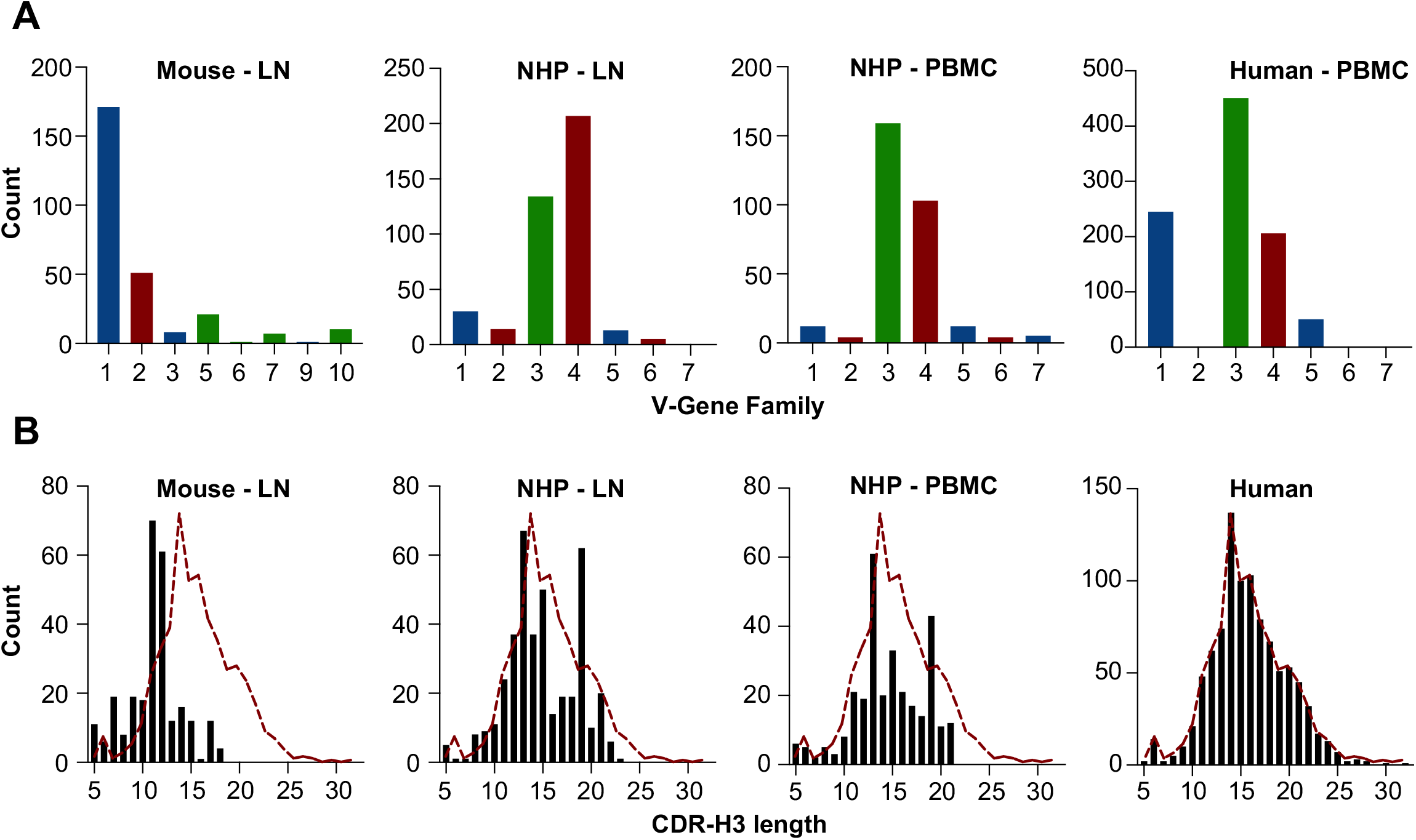
**(A**) Distribution of V-gene family utilisation or (**B**) CDHR-H3 lengths of B cell receptor sequences recovered from: GC B cells (B220^+^IgD^-^GL7^+^) within the draining iliac LN of C57BL/6 mice (n=3) 14 days after immunisation with S; RBD- and S-specific B cells (CD20^+^IgD^-^IgG^+^) in the the draining iliac LN or PBMC of a single macaque 14 days after a second immunisation with S; or RBD- and S-specific B cells (CD19^+^IgD^-^IgG^+^) within PBMC of convalescent COVID-19 subjects (n=6; as reported in Juno et al., 2020)

